# Novelty at second glance: A critical appraisal of the novel-object paradigm based on meta-analysis

**DOI:** 10.1101/2020.12.10.419457

**Authors:** E. Takola, E. T. Krause, C. Müller, H. Schielzeth

## Abstract

The study of consistent individual differences in behaviour has become an important focus in research on animal behaviour. Behavioural phenotypes are typically measured through standardized testing paradigms and one frequently used paradigm is the novel object test. In novel object tests, animals are exposed to new (unknown) objects and their reaction is quantified. When repeating trials to assess the temporal consistency of individual differences, researchers face the dilemma of whether to use the same or different ‘novel’ objects, since the same stimulus can result in habituation, while exposure to different objects can result in context-dependent responses. We performed a quantitative assessment of 254 effect sizes from 113 studies on novel-object trials to evaluate the properties of this testing paradigm, in particular the effect of object novelty and time interval between novel-object trials on estimates of individual consistency. We found an increase of sample sizes and an increase of estimates of repeatabilities with time. The vast majority of short-term studies (<one month) used different novel objects, while long-term studies (>one month) used either the same or different novel objects about equally often. The average estimate for individual consistency was r = 0.47 (short-term r = 0.52, long-term r = 0.44). Novelty, time interval between trials and their interaction together explained only 3% of the total heterogeneity. Overall, novel-object trials reliably estimate individual differences in behaviour, but results were very heterogeneous even within the same study species, suggesting susceptibility to unknown details in testing conditions. Most studies that measure novel-object responses in association with food label the trait as neophobia, while novel-object trials in a neutral context are labelled variously as boldness/shyness, exploratory behaviour or neophobia/neophilia. Neophobia/neophilia is also the term most specific to novel object presentations. To avoid ambiguity, we suggest object neophobia/neophilia as the most specific label for novel-object responses.

## INTRODUCTION

Consistent individual differences in behaviour are widespread in nature. For a long time, individual differences were considered noise around an optimum niche value (Parker & Smith, 1990; Carere & Eens, 2005; Wolf & Weissing, 2012). Nowadays it is increasingly appreciated that intraspecific variation is of widespread adaptive importance and can affect reproductive success (Smith & Blumstein, 2008), growth rates (Royauté, Berdal, Garrison, & Dochtermann, 2018), metabolic rates (Holtmann, Lagisz, & Nakagawa, 2016) and even population dynamics (Levin, Tolimieri, Nicklin, & Sale, 2000). Is has also been established that individual differences in behaviour often have a heritable basis (Dingemanse & Réale, 2005; Schielzeth, Bolund, Kempenaers, & Forstmeier, 2011) and can thus respond to selection. The study of individual differences has therefore become an important topic in behavioural ecology.

Particularly interesting are individual differences in context-general responses to environmental challenges. They can, for example, explain maladaptive behaviour in some contexts if the same behaviour is advantageous in other contexts (Sih, Bell, Johnson, & Ziemba, 2004). Individual differences in responses across contexts are variously called animal personalities, behavioural syndromes, coping styles or temperament (Réale, Reader, Sol, McDougall, & Dingemanse, 2007). Apart from contextual consistency, another hallmark of animal personality is that traits are individually stable over time (Sih et al., 2004; Kaiser & Müller, 2021). Most studies on animal personality use standardized experimental setups with repeated measurements per individual to estimate contextual and temporal consistency.

The degree of individual consistency is typically measured either by correlations (Pearson’s and Spearman’s or Kendall’s) or by repeatability among repeated observations of the same individuals (Bell, Hankison, & Laskowski, 2009). Repeatability is expressed as the ratio of between-individual variance to the total phenotypic variance of the population (Sokal & Rohlf, 1995), thereby generalizing pairwise correlations to multiple measurements, and is hence also known as an intra-class correlation coefficient (ICC) (Nakagawa & Schielzeth, 2010). Repeatabilities can be estimated from mixed-effects models that offer substantial flexibility in controlling for confounding random and/or fixed effects (Réale et al., 2007). Repeatabilities that control for confounding effects are often larger than unadjusted repeatabilities of the raw data and are known as adjusted repeatabilities (Nakagawa & Schielzeth, 2010).

The assessment of behavioural phenotypes can be achieved through various experimental procedures. One popular method is novel-object presentations (Yerkes & Yerkes, 1936). In novel-object presentations, animals encounter an item that they had never seen before (thus a novel object) and their behavioural responses are quantified, particularly the amount of movement, approach distances or approach latencies (Yerkes & Yerkes, 1936; Greenberg, 1990; Guenther & Brust, 2017). There are multiple variants of novel-object trials and one area of controversy is how the resultant behaviours are best labelled. Another question is how a response to novelty can be measured repeatedly to establish that behavioural scores represent individual differences rather than within-individual phenotypic flexibility.

Behavioural responses in novel-object trials are mostly used to measure shyness-boldness, exploration or neophilia/neophobia. Novel-object trials are not the only testing paradigm to measure these traits. Shyness-boldness is also often quantified by startle response trials, emergence from shelter, response to predator or by mirror image trials (Ioannou, Payne, & Krause, 2008; Noer, Needham, Wiese, Balsby, & Dabelsteen, 2015). Réale et al. (2007) suggest that shyness and boldness refer to an individual’s reaction towards a risky situation in general, but not to novelty *per se*. Exploratory behaviour is also often quantified by open field or novel-environment trials. Réale et al. (2007) propose the gradient of exploration-avoidance for the behavioural response to a new situation in general, including novel environments and novel objects. Neophilia/neophobia is a more specific term to novel-object trials, although neophilia/neophobia is sometimes also used for novel environment trials (Greggor, Thornton, & Clayton, 2015). Mettke-Hofmann (2012) therefore distinguishes object neophilia/neophobia for novel-object trials from spatial neophilia/neophobia for novel-environment trials.

Independent of the question of labelling is the question of repeated presentations and how they shall be best conducted in the experimental design. While the first presentation of a novel object can generate the intended response, upon second presentation of the identical item objects are no longer novel. Thus, the second presentation may trigger a different response, because repeated exposure to the same stimulus can lead to a reduced strength of the behavioural response (Berlyne, 1966). The alternative is to use different unknown objects, which might trigger different responses, for example, if they differ in conspicuousness or perceived riskiness. Greggor, Jolles, Thornton, and Clayton (2016) suggest that objects should be used that differ slightly but clearly. However, similarity and differences are ambiguous categories and what might be perceived as similar by some might be seen as different by other individuals. Furthermore, some species might habituate to novel stimuli *per se* (Réale et al., 2007), such that even slightly different novel objects do not trigger the same behavioural response upon second presentation.

Moreover, the effect of using the same or different objects in repeated trials likely depends on the time interval between repeated presentations. The degree of novelty in these repeated trials is certainly the result of perception and memory (Greggor et al., 2016), but our understanding of animal memory and cognition mechanisms is still incomplete, in particular when it comes to a large range of taxa. It is likely that the effects of use of novel objects differ between short-term replication (within hours, days or weeks) and long-term replication (after months or years). Therefore, the temporal interval between trials should be taken into account when studying behavioural consistency.

We here review the results of novel-object trials using meta-analytic techniques (Koricheva, Gurevitch, & Mengersen, 2013; Gurevitch, Koricheva, Nakagawa, & Stewart, 2018). Metaanalysis is a powerful tool for research synthesis in science, as it provides an objective and replicable quantitative overview of literature. Importantly, the use of moderators as fixed effects allows for the identification of context-dependencies. Although a common criticism of meta-analytical methods highlights the pooling of incomparable effect sizes (also known as ‘apples and oranges problem’), we addressed the issue of diverse study designs by adding variables as moderators and accounting for multi-level variation. Moreover, we accounted for phylogenetic correlations, since closely related species might react similarly to the same stimuli (Nakagawa & Santos, 2012). Thus, we were able to explore the impact of various effects on the consistency of behavioural traits from multiple studies.

The aim of our meta-analysis was to examine whether the use of same or different objects during repeated novel-object tests is affecting the temporal consistency of individual responses. Specifically, we conducted a meta-analysis to estimate the effect of time interval between repeats, novelty and their interaction. Apart from time interval and novelty, we used a variety of moderators, such as context of testing and domestication level of animals, to account for variation in estimates of temporal consistencies. In addition, we summarize and discuss variation in terminology when labelling response behaviours and present an overview of the most common traits used as proxies to describe different behavioural phenotypes.

## MATERIALS AND METHODS

We used systematic reviewing techniques to evaluate the properties of the novel-object paradigm for quantifying consistent individual difference in behaviour (Koricheva et al., 2013). Our methodology followed the PRISMA protocol (Moher, Liberati, Tetzlaff, Altman, & PRISMA Group, 2009).

### Data Collection

We conducted a search in the Web of Science (WoS) Core Collection 5.24. The query included not only the term novel object (novel object*), but also words related to behavioural phenotypes (e.g. neophob*, neophil*, bold*, shy*) and the time range was set to 1990-2020 (Supplement S1). The early 1990s were the time when novel-object trials were first used systematically to quantify individual differences for context-general behavioural traits (Greenberg, 1990), such that we are confident that our search range includes all relevant studies. We also searched for the search term explorat*, but the number of hits was very large (more than 3,000 additional publications) and since the label exploration is primarily used for other behavioural assays, we did not include the term in the final search. The WoS Category was limited to Behavioural Sciences and duplicates were removed, resulting to 3,599 publications that were used for more detailed screening. The literature search was finalized on the 15th July 2020.

### Inclusion and Exclusion Criteria

We searched for empirical studies that used novel-object presentations and quantified the responses of individual animals to these objects. A novel object should be unfamiliar to focal individuals (at least upon first presentation). A novel object should also be unusual to focal animals so that we do not expect an evolved attraction to these objects, thus excluding objects that represent natural resources of a species. We did include novel food sources in our analysis if the novel food was sufficiently different from the natural food of a species. This included studies that use artificial dyes to stain natural food if the novel food colour was considered sufficiently novel and unusual.

We screened studies based on the following criteria (Supplement Table S1). First, studies should be done with outbred, non-human animals with unimpaired physical condition. Second, studies should use a novel-object paradigm, thus excluding mirror-images, live conspecifics, taxidermy mount presentations and natural resources of a species. Third, studies should have repeated novel object trials at the level of individuals. Fourth, studies should report relevant correlations or repeatability as a measure of individual consistency.

We conducted the screening process in two stages. We first screened title and abstracts, which excluded 2,928 publications, mostly because they did not represent empirical studies, they were done on humans, they did not use systematic novel-object presentations or they did not study individual differences (Figure 1). Only clearly non-fitting cases were excluded during abstract screening and doubtful cases were taken forward to the next step. We next screened full-texts of the remaining 671 publications. Screening of full-texts was done independently by two people (ET and HS) and conflicts were resolved jointly. Full-text screening was focused on the same general criteria and also on whether relevant effect and sample sizes were reported. Another 558 publications were excluded during full-text screening (Figure 1). One study was opportunistically added to the final dataset. Consequently, 113 studies matched our inclusion criteria and generated 254 effect sizes.

**Figure 1.**
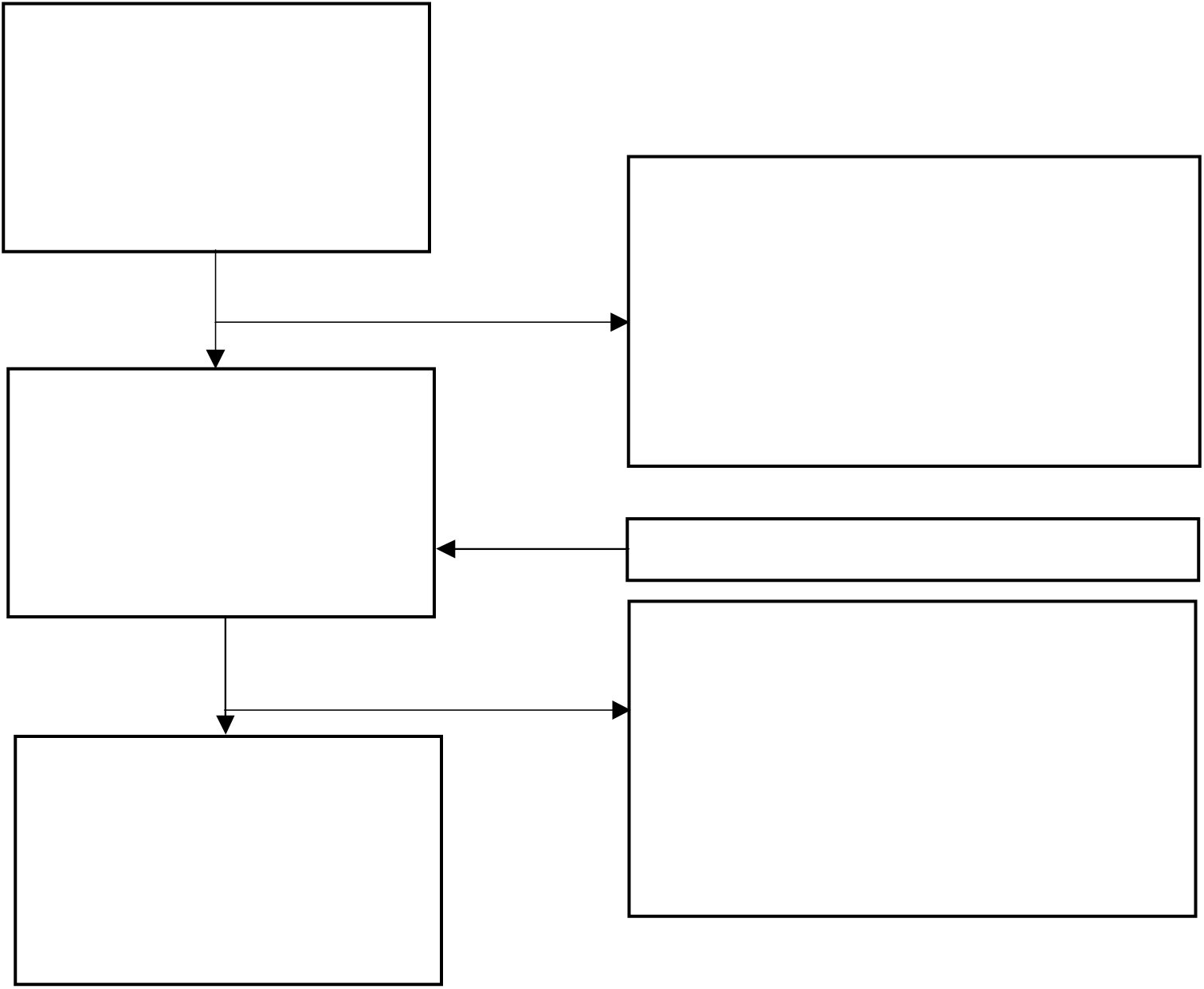
PRISMA diagram with abstract and full text screening results. Numbers show the number of publications that were excluded or included.

### Data Extraction

We extracted pairwise correlation coefficients (Pearson’s, Spearman’s or Kendall’s) and repeatabilities (R or ICC), as we were interested in the temporal consistency of behavioural responses between trials. Effect size typically corresponded to two rounds of novel-object trials with the same set of individuals. In cases with more than two rounds of testing or when multiple responses were quantified, multiple effect sizes were extracted from one study. When combined repeatabilities were reported for more than two trials, we extracted these repeatabilities as the relevant effect sizes. In five cases correlation measures were extracted from graphs using the metaDigitise package, version 1.0.1 (Pick, Nakagawa, & Noble, 2019).

For each effect size we extracted information related to (i) publication (year, authors, and journal), (ii) animals tested (species, sample size, and domestication status), (iii) testing conditions (novelty of the object in the repeated trials, time interval between trials, and context of testing), (iv) response behaviour being quantified (specific individual behaviours, response type, see below), (v) analyses being conducted (whether multiple personality traits were assayed, whether repeatabilities were calculated from non-Gaussian generalized linear models) and (vi) the terms used to describe the behavioural phenotype (see Supplement Tables S2 and S3 for a detailed description).

The novelty of the object in repeated trials was a parameter of key interest in our analysis. When the novel objects were the same but of different colours, we considered them as different objects. Context of testing was categorised into (i) neutral context, (ii) novel object next to food or (iii) novel object next to nest. For domestication status we distinguished between (i) domesticated, (ii) lab-reared, (iii) wild-caught and lab-tested, (iv) wild and tested in the wild (Mathot, Dingemanse, & Nakagawa, 2019). For response type and behaviour we recorded the specific trait being quantified (if it was a single behavioural response), whether the response was a composite of multiple behaviours within the same trial (often principle component scores of multiple behaviours scored within the same trial or other synthetic response scores based on multiple components of behaviour) or whether the response was an average calculated across multiple (sub)trials. We did not record transformations being used, since we consider this a decision of individual researchers to best quantify the behaviour, similar to the researcher’s decision to record a specific response behaviour and not another. For the same reason, we also did not distinguish between parametric (Pearson) and nonparametric (Spearman or Kendall) correlations. However, a few studies analysed behavioural phenotypes as binary responses or using Poisson models and these might produce systematically lower consistency measures and were therefore recorded.

The time interval was recorded in days, assuming 30 days in a month and 365 days in a year when converting from descriptions in publications). Since our dataset included many species with different life-histories, we also tried to standardize time intervals by dividing them with the species’ lifespan (compiled from the AnAge database) (Tacutu et al., 2018) to express the time interval as a proportion of lifespan. However, raw time interval measures and lifetime standardized measures were highly correlated, r = 0.94 (and results were qualitatively unaffected), such that we used log-transformed time interval in days as a moderator in our analysis.

### Effect size and weighing in meta-analytic models

We used R 3.6.3 for all analyses (R Core Team 2020). Correlation and repeatability measures were transformed using Fisher’s Z-transformation as implemented in the *escalc* function of the *metafor* package, ver. 2.4.0 (Viechtbauer, 2010). Effect sizes were weighted by the inverse of sampling variance in all analyses.

### Meta-analyses and meta-regressions

We conducted a phylogenetic multilevel meta-analysis in order to estimate the overall effect. Phylogenetic information was downloaded from Open Tree of Life version ott3.2 (Hinchliff et al., 2015) using the *rotl* (ver. 3.0.11) R package (Michonneau, Brown, & Winter, 2016). After constructing an ultrametric phylogenetic tree (Supplement Fig. S1) using the Grafen method (Grafen, 1989), we converted the tree to a correlation matrix. This matrix was fitted as a random effect in our meta-analytic model, along with random effects for effect size ID, study ID, and species ID. The analysis was performed first using the complete dataset and then separately for major taxonomic groups (mammals, birds, fish, reptiles, and insects). Weighted random-effect-only meta-analytic models were fitted using the *rma* function of the *metafor* package.

Besides the random-effect-only meta-analytic model, we also fitted a meta-regression with moderators (Table S3), once for the complete dataset and once for every major taxonomic group represented by more than ten publications in our dataset (mammals and birds). As moderators we fitted the time interval between repeated tests (log-transformed), novelty (2 levels), domestication status (4 levels), correlation type (2 levels), a binary indicator for non-Gaussian linear models, a binary indicator of whether multiple behavioural tests were performed in the study (other than the novel object), response type (3 levels), testing context (3 levels) and the interaction of novelty with time interval. As above, the random effects of the meta-regression were the effect size ID, study ID, phylogeny and species.

Heterogeneity (*I^2^*) was examined for multiple levels in every model in our meta-analysis, including the subsets of different clades (Nakagawa & Santos, 2012). We also calculated marginal *R^2^* to estimate the proportion of variance explained by fixed effects (Nakagawa & Schielzeth, 2013). The variance explained by individual predictors were calculated by fitting a specific predictor in a meta-regression, followed by calculation of marginal *R*^2^.

### Sensitivity analyses

We conducted influence diagnostics and sensitivity analyses to evaluate the robustness of our results. For the influence diagnostics we used the *influence* function of the *metafor* package, ver. 2.4.0 (Viechtbauer, 2010), in order to identify influential studies using Cook’s distance and *rstudent* test. The diagnostics showed five potential outliers in the dataset (Supplement Fig. S2). We therefore refitted the meta-analytic model again while excluding the five influential effect sizes. Since the overall estimate was not significantly affected, we present the analysis of the full dataset.

### Publication bias

We tested for publication bias qualitatively through visual inspection of funnel plots and quantitatively by Egger’s regression (Egger, Smith, Schneider, & Minder, 1997). Funnel plots were generated by plotting effect sizes against inverse sampling variance and inverse standard error. Egger’s regression estimates funnel plot asymmetry as an indicator of publication bias. In addition, we examined the possibility of time-lag bias, which is the decrease of effect sizes with time (Trikalinos & Ioannidis, 2006). The test for differences in effect sizes between studies that used novel-object trials as the only personality scoring paradigm versus studies that used multiple measures of personality traits also served as a test for publications bias. We expect studies with a single behavioural measure to be more likely to report statistically significant temporal consistency than studies that report on multiple behavioural traits, of which only a subset might be significantly repeatable.

## RESULTS

Screening of 3,599 abstracts and full texts resulted in 220 studies that used novel-object trials to quantify individual behaviour in non-human animals. 163 (74%) of these studies replicated novel-object trials for all or for a subset of individuals. After excluding 50 studies with repeated novel-object trials that did not allow an extraction of effect sizes for temporal consistency, we found 254 effect sizes from 113 studies (Supplement Table S6) to be included in the analyses. This dataset encompassed 69 species (21 mammal, 35 bird, 5 fish, 4 reptile and 4 insect species) (Supplement Fig. S1).

### Testing practices

Sample size ranged from 5 to 567 individuals per effect size estimate (average ± SD: 48.7 ± 58.6) and sample size increased significantly by about 2.5% per year (effect of year of publication on log(N) sample size: b = 0.026 ± 0.009, t_192_ = 2.12, *p* = 0.002, Figure 2).

**Figure 2.**
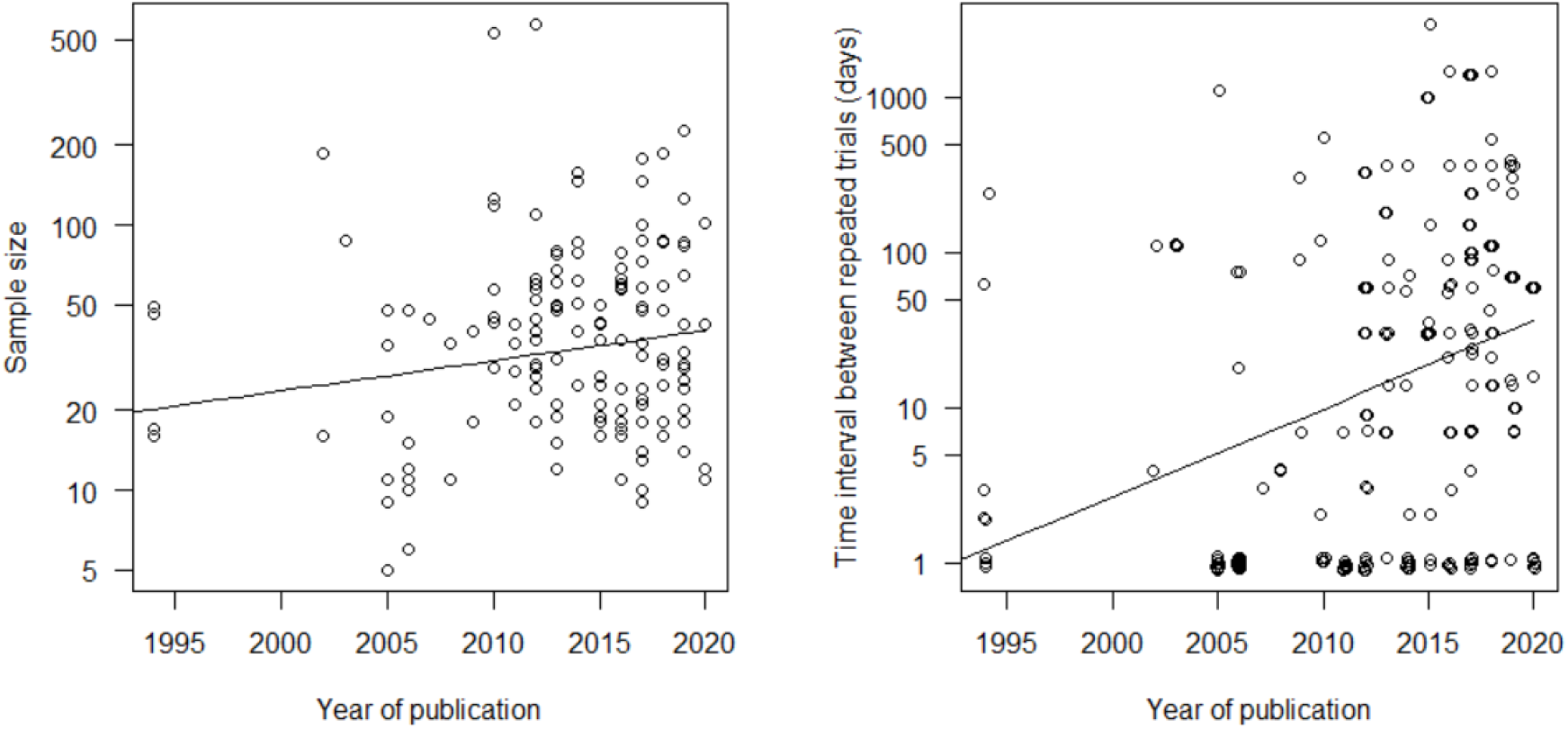
Temporal trends of sample size and time interval between repeated trials. Open dots show effect size estimates by sample size and time interval. The sample size and time interval are shown on a log scale.

The time interval between two consecutive trials ranged between a few hours and four years (<0.1% to 82% when expressed relative to the expected lifespan of the focal species). Sixty-five effects sizes (32%) refer to trials repeated on the same (5 effect sizes) or on consecutive days (64 effect sizes). 62% of the effect sizes were calculated from replications after at least one week, 42% after more than one month and 8% after at least one year. Studies over longer time periods became more popular over the years with an increase in the time interval between trials of about 13% per year (effect of year of publication on log(time interval): b = 0.129 ± 0.022, t_192_ = 5.86, P < 10^-5^, Figure 2). In the following, we operationally define effect sizes calculated from repeats less than one month apart as short-term replications and those with longer intervals as long-term replications.

Seventy-four studies used different objects in repeated trials, 33 studies used the same objects and six studies used both. Most short-term studies (83% of effect sizes for short-term repeatabilities) used different objects, while same ‘novel’ objects where used more often when addressing long-term consistencies (only 32% different objects among estimates for long-term repeatabilities, Supplement Table S2).

Seventy-five studies conducted novel-object presentations in a neutral context (73% of effect sizes), 30 studies next to a food source (21% of effect sizes) and 8 studies inside or close to nest (6% of effect sizes). Most studies calculated individual consistencies for a specific response behaviour (75% of effect sizes), some used principle component or other composite scores calculated from multiple behavioural components measured in the same trial (13% of effect sizes) or calculated individual temporal consistencies after averaging across multiple trials (12% of effect sizes). Most studies (84%) used novel-object trials along with other standardized personality assays (such as open field trials, startle responses or intruder trials), while only 15 studies (16%) focused on the behavioural consistency for novel-object trials only (Supplement Table S2).

### Overall effect sizes and heterogeneities

The overall effect of the phylogenetic meta-analysis was strong and significantly greater than zero (β_0_ = 0.52, CI = [0.46, 0.58]), which is equivalent to a correlation of r = 0.47. Heterogeneity among effect sizes was high (*I^2^_total_* = 81%). Variation among studies and among effect sizes accounted for 54% and 26% of this heterogeneity (Table 2), respectively, while species identity and phylogenetic relationships explained a negligible part. The average short term repeatability was r = 0.52 (r = 0.51 for time intervals of up to one week and r = 0.59 for time intervals between one week and one month, Figure 3), while the average long term repeatability was r = 0.40 (r = 0.41 for time intervals of one month to one year and r = 0.33 for time intervals of more than one year).

**Figure 3.**
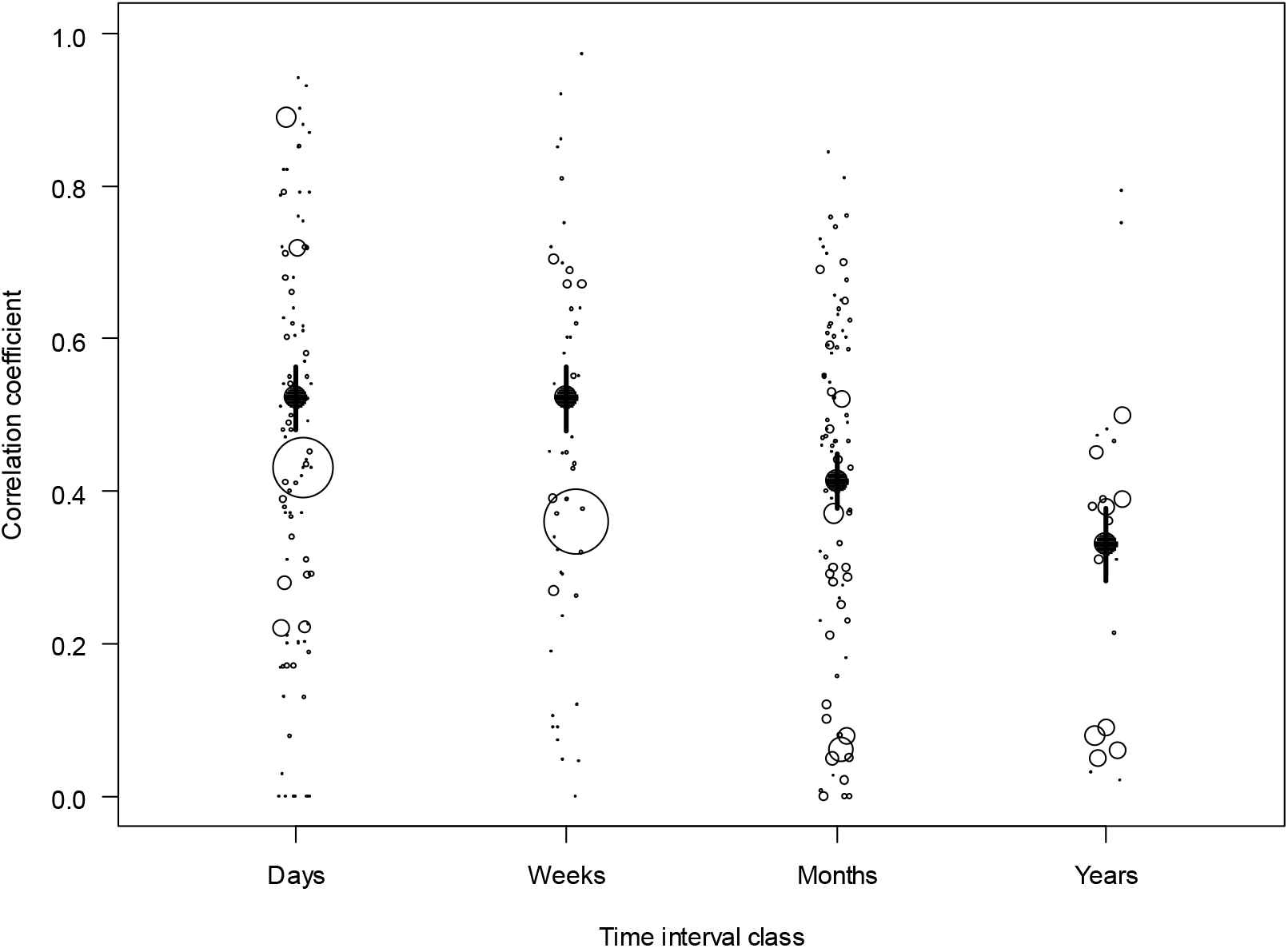
Overall effect sizes for four classes of time intervals between repeated trials. Open dots show effect sizes (dot size scaled by sample size) and black dots and bars show random-effect-only meta-analytic model estimates ± SE. Days = 0-6 days, Weeks = 7-30 days, Months = 31-364 days, Years = 365+ days.

**Table 1.** Summaries and results from phylogenetic multilevel meta-analyses. LCI and UCI indicate the lower and upper limits of the 95% confidence intervals.

| | $N_{\text{E.S.}}$ | $N_{\text{studies}}$ | $N_{\text{species}}$ | Zr | SE | LCI | UCI | z | $p$ | r |
| --- | --- | --- | --- | --- | --- | --- | --- | --- | --- | --- |
| Overall | 254 | 113 | 69 | 0.52 | 0.03 | 0.46 | 0.58 | 17 | <.0001 | 0.47 |
| Mammals | 82 | 34 | 21 | 0.54 | 0.06 | 0.42 | 0.65 | 9.14 | <.0001 | 0.49 |
| Birds | 141 | 62 | 35 | 0.54 | 0.042 | 0.46 | 0.62 | 12.75 | <.0001 | 0.49 |
| Fish | 19 | 10 | 5 | 0.52 | 0.15 | 0.22 | 0.81 | 3.44 | 0.0006 | 0.46 |
| Reptiles | 8 | 3 | 4 | 0.07 | 0.05 | -0.02 | 0.17 | 1.49 | 0.17 | 0.074 |
| Insects | 4 | 4 | 4 | 0.43 | 0.07 | 0.28 | 0.58 | 5.7 | <.0001 | 0.40 |

**Table 2.**
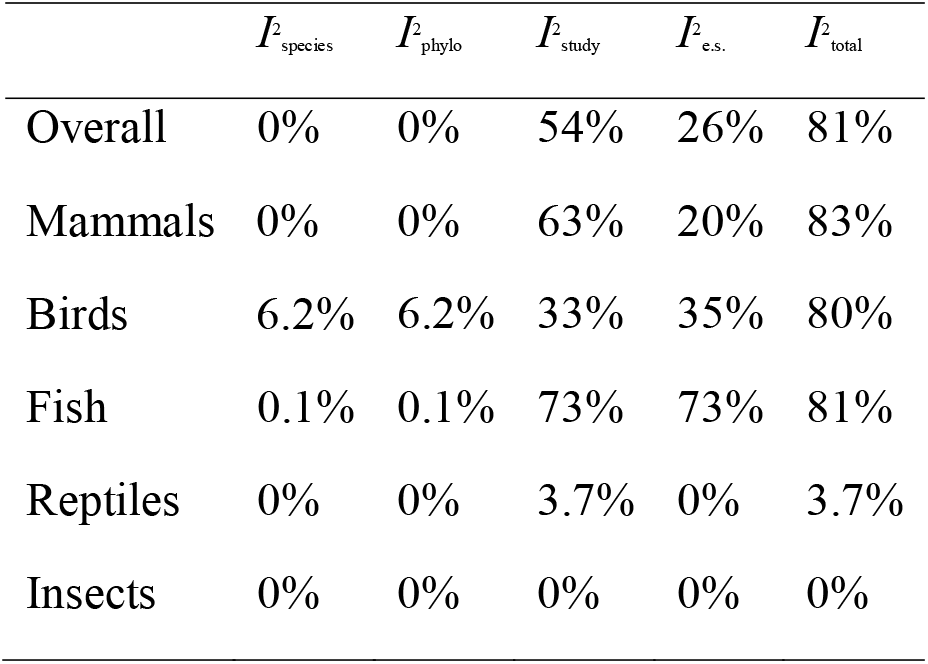
Total heterogeneity in effect sizes across hierarchical levels of random effects for the overall dataset and for subsets of the data. Accuracy to one decimal only for effects <10%.

We repeated the analysis separately for the subsets of mammals, birds, fish, reptiles and insects. Mammals, bird, fish and insects showed strong and significant consistencies of behaviour (all r > 0.40), while individual consistency was low and non-significant for reptiles (r = 0.074, Table 1). Total heterogeneity was particularly high in the subsets of mammals, birds and fish (all *I^2^_total_* > 80%) but not for insects and reptiles (*I^2^_total_* < 4%) (Table 2). Between-study heterogeneity was particularly high for the subset of mammals and fish (*I^2^_study_* > 63%), moderate for birds (*I^2^_study_* = 33%) and low for insects and reptiles (Table 2).

### The impact of novelty in repeated trials and time interval between repeats

The amount of total heterogeneity in overall effect indicated scope for effects of moderators. We therefore fitted a meta-regression with novelty, time and their interaction as moderators. These moderators explained 3% of the variance and did not have a significant effect on the correlation (*Q_M_* = 4.21, *p* = 0.20). Novelty had a low and non-significant effect on behavioural consistency and as expected, time yielded a negative estimate (shorter time intervals resulted in higher repeatability estimates). The estimate for the interaction was negative (the effect of time interval was stronger if objects were different), but not significantly different from zero (β_int_ = −0.002, CI = [-0.0485, 0.0440], *p* = 0.92). Similar trends were observed in the subsets of mammals and birds (Figure 4). In the overall model and the subset of birds, these moderators explained less than 4%, but in the subset of mammals, they explained 6%. Even though meta-regression did not show a significant effect of time, long-term consistencies seems to be marked lower than short-term consistencies when the data are broken down to time interval classes (Figure 3).

**Figure 4.**
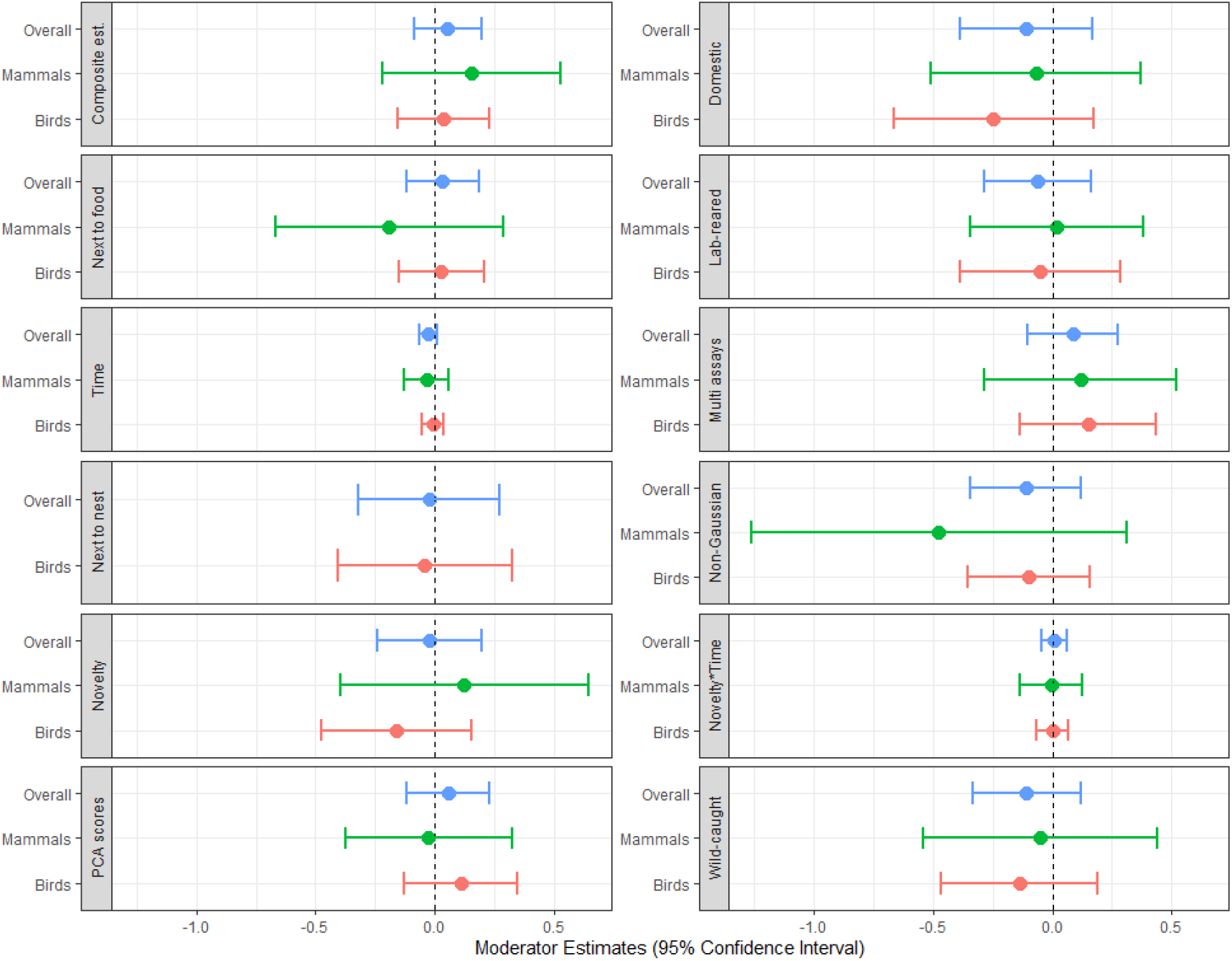
Forest plot showing the results of meta-regressions using the full dataset and subsets of bird and mammal species. Moderators were time interval, novelty, the interaction of novelty and time, the domestication gradient of testing and testing context (position of novel object relative to resources). The reference category combination in the model was the same object, wild-caught animals tested in the wild, neutral context, Gaussian models, single behaviour, repeatability estimate and only novel objects as the only personality trait being assayed.

### The impact of other moderators

We explored effects of additional moderators in the meta-regression model by fitting each one of them in a meta-regression model. Domestication status accounted for a low fraction of variance (*R^2^_dom_* = 1%) and was not significantly correlated with the overall effect size. In the subset of mammals, domestication status explained 2% of variation and for birds 1%. However, the levels of domestication status did not show consistent estimates across different subsets of the data (Figure 4). Testing context explained only 1% of the total heterogeneity. The type of response (single behaviours, aggregates of multiple components and averages across trials) had no significant effect and the effect of estimation by non-Gaussian models was also non-significant. All moderators explained less than 4% in all cases, only in the subset of mammals the response type explained 8% and non-Gaussian GLMMs explained 6%.

### Publication bias and sensitivity analysis

For sensitivity analysis, we refitted the overall meta-analytic model without five particularly influential studies (Supplement Fig. S2). The model estimate marginally decreased from 0.52 to 0.49 (CI = [0.44, 0.54]), whereas the total heterogeneity dropped from 81% to 73%.

Visual inspection of the funnel plot showed only weak asymmetry of effect sizes (Figure 6). However, Egger’s test identified significant asymmetry (t_199_ = 3.56, *p* = 0.0004) and a subsequently trim-and-fill method estimated 15 missing effect sizes. Hence, our results for the overall effect might be slightly biased upwards. We tested for time-lag bias by fitting a meta-regression with publication year as a predictor. The slope showed a negative trend (β = −0.01 (CI = [-0.02, 0.0009], *Q_M_* = 3.23, *p* = 0.072) and explained 4.1% of variance. Studies that report multiple behavioural traits had non-significantly larger consistency estimates than studies that focus on novel-object trials. This result is not indicative of publication bias.

### Reproducibility within species

The amount of heterogeneity explained by species was estimated to zero in the overall metaanalysis. However, most species were used only in one or few studies. Three species, however, were the focal species of more than three studies and we inspected the consistency of estimates within these three species (guinea pig *Cavia aperea*, zebra finch *Taeniopygia guttata* and great tit *Parus major*) more closely. Estimates of individual consistency in response to novelty of the guinea pig were done with two laboratory populations (domestic guinea pigs, and wild-derived cavies) in the same research lab, all with the novel object in a neutral context and they used either latency to approach or the number of touches as a response. Nevertheless, estimates varied widely (Figure 5). Estimates with zebra finches were all done in seven different outbred laboratory populations (including the study with the second largest sample size in our dataset) and were either performed in a neutral context or close to food. Estimates varied widely (Figure 5) within contexts and even within the same population with multiple estimates. Estimates for the great tit were particularly heterogeneous in context (neutral, near food or near nest) and they were conducted in the wild, in the lab with wild-caught birds or with lab-bred individuals. However, the scatter of estimates was similar to the cases of guinea pigs and zebra finches (Figure 5).

**Figure 5.**
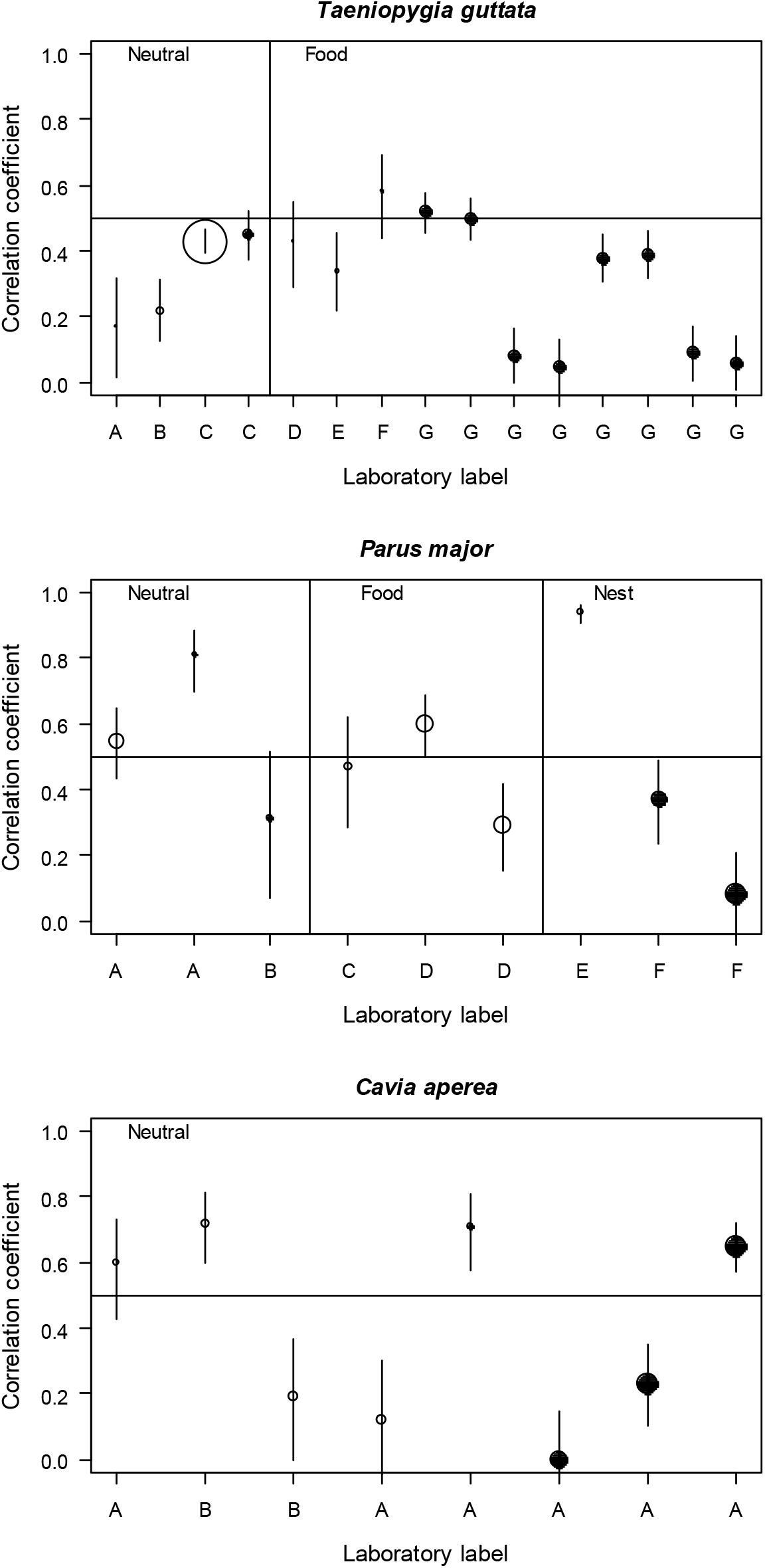
Repeatability (estimates ± SE) of behaviour for the most popular species in our dataset: zebra finch (*Taeniopygia guttata*), great tit (*Parus major*) and guinea pigs (*Cavia aperea*). Open and filled dots are used to indicate short and long time intervals respectively. The size of the dots is scaled by sample size. Different letters for the laboratory label mark different populations of animals.

**Figure 6.**
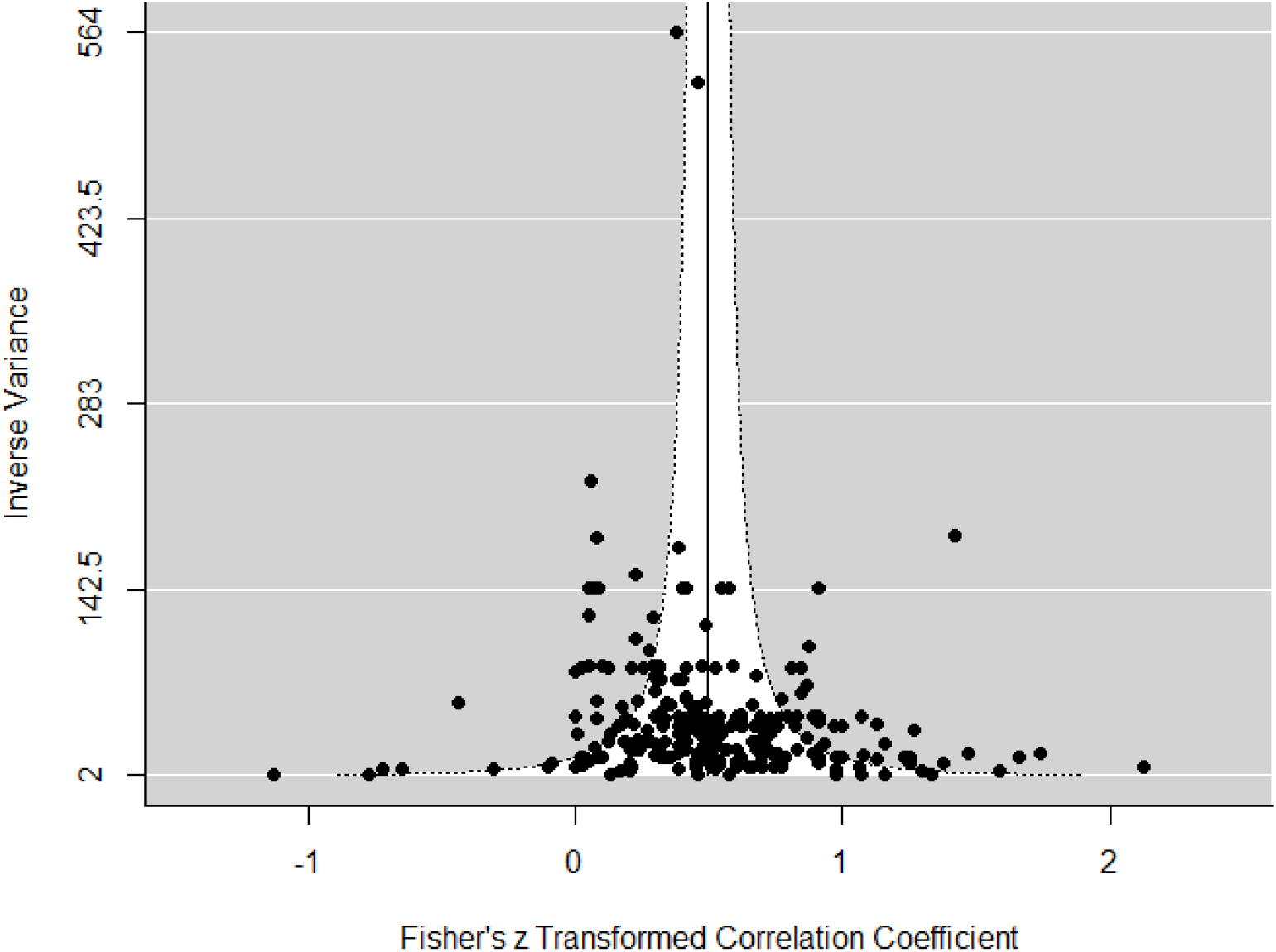
Funnel plot of inverse sampling variation against effects sizes for an assessment of publication bias.

### Terminology

Most studies labelled responses to novel objects as either neophobia/neophilia (48 studies, 42%), shyness/boldness (31 studies, 27%) or exploration (22 studies, 19%), while more rarely occurring labels were fearfulness (5 studies), approach-avoidance (2), risk-responsiveness (2) and activity (1) (Supplement Tables S4, S5). Eight studies did not use any general labels for the traits being measured. Labelling was associated with testing context, with an even stronger bias towards neophobia when novel-objects were place next to food or nests (70% across these two contexts) and a more even distribution across neophobia/neophilia, boldness/shyness and exploration when the novel object was located in a neutral place (Supplement Table S4, Fig. S3).

## DISCUSSION

Our meta-analysis of measures of individual temporal consistencies quantified in novel object trials revealed an overall strong and significant repeatability of responses to novel objects (r = 0.47). This estimate is substantially larger than an estimate of average repeatability in behaviour (r = 0.37; (Bell et al., 2009), which demonstrates that the novel-object paradigm is a useful and reliable way to quantify consistent individual differences among individuals. It had been predicted that the repeatability declines with an increase of the time interval between observations (Bell et al., 2009), a trend that we can visualize in our analysis. Any state-dependent causes of individual differences are likely to be temporally autocorrelated, such that short-term repeatabilities should be higher than long-term repeatabilities. Environmental variables also tend to be temporally autocorrelated, which can lead to pseudo-repeatability, in particular when individuals select their different microenvironments or individualized niches (resulting in ‘recurrent environments’) (Dupré, 2014). Both state-dependent and environment-dependent autocorrelation are predicted to lead to larger shortterm repeatabilities.

### General evaluation of the testing paradigm

The rather high overall repeatability in response to novel objects shows that novel object trials provide generally suitable assays for a quantification of temporal consistencies and thus also differences among individuals. However, we found substantial heterogeneity in effect sizes, though mostly between studies and not between species. The large heterogeneity poses the question of whether differences between studies reflect genuine differences between populations or whether they reflect differences in the uncontrolled aspects of the experimental setup. There are many reasons, why populations may differ in the relative magnitude of individual differences. For example, populations might have been exposed to different selective regimes, such as urban versus rural populations (Miranda, Schielzeth, Sonntag, & Partecke, 2013) or captive versus wild populations (Herborn et al., 2010). In addition, population size might affect the amount of standing genetic variation and thus the phenotypic variation for behavioural traits. Moreover, the environment might affect the magnitude of state-dependent individual differences (Sih et al., 2015), which might arguable be larger in the wild than in captivity, though empirical evidence is scarce. Any such differences in population background, population size and the magnitude of state variation could give raise to heterogeneity in effect sizes.

Hence, heterogeneity might well be real and relevant to understand variation in individual behavioural traits related to personality. However, it is also important to consider the nonexclusive alternative, that experimental setups of novel object trials differ in how reliably they capture individual differences. This is an important concern, since most studies used response to novelty as a trait to be correlated with other behaviours (Guenther & Brust, 2017), endocrine measurements (Arnold et al., 2016) or reproductive success (Collins, Hatch, Elliott, & Jacobs, 2019) and these relationships might be systematically underestimated, if behavioural measurements contain substantial measurement error. For example, experimental setups might trigger different responses depending on short-term state fluctuation (e.g. the state of hunger). Furthermore, we usually know far too little about which objects might trigger sufficient interest in animals and which objects are perceived as intimidating, which likely is influenced by size, colour, shape and odour of the object as well as familiarity to similar-looking, known objects. Some of the heterogeneity might not represent differences in behaviour among individuals, but rather variation in novel-object trials themselves, thus potentially impairing robustness of the paradigm. Two lines of evidence suggest that there are some issues with the experimental setup.

First, under the premise that novel-object trials are designed to measure context-general personality traits, we would expect consistent findings at least within species. However, the between-species component of heterogeneity was very low and replicate studies within three specific species (guinea pig, zebra finch and great tit) show substantial differences in estimates (Figure 5). It could be argued that these reflect genuine population differences in the case of the zebra finch and great tit, but this seems unlikely in the case of the guinea pig, since all studies on this species were performed in the same laboratory (with two different populations of animals).

Second, most moderators showed no significant effect, although at least some of them had well-justified predictions. For example, the effect of averaging is bound to have a systematic effect on repeatabilities. In particular, behavioural scores that are averaged across multiple trials should have higher repeatabilities, because environmental noise is averaged out (Nakagawa & Schielzeth, 2010). We had therefore predicted that averaging within trials (e.g. by using principle component scores) and even more averaging across trials increases the estimates. However, this effect was not significant. Lack of significance for such moderations suggests an overall large amount of noise in the data.

### Specific design decisions

Selection of novel objects is very important, as it can induce different reactions (Greggor et al., 2015). Interestingly, experimental design decision such as the use of the same or different novel objects for the test replications seems to play a very minor role in influencing the magnitude of individual differences, since on average estimates were not significantly different. However, the vast majority of short-term repeatability estimates was based on the use of different novel objects. This is a useful decision for the test setup for two reasons. First, shorter time intervals will make it more likely that individuals remember specific objects (Bell et al., 2009). Second, novel-object trials are intended to quantify context-general aspects of behaviour, hence it is the repeatable component in response to different objects that matters in most cases. Over extended time periods, however, it seems less likely that individuals actually remember a specific encounter. Indeed, about half of the long-term studies that used the same novel objects when re-testing were done months or years after a first encounter and this test design had no systematic effect on the magnitude of consistent individual differences, suggesting that the quantified behaviours are as comparable as with different novel objects.

The phylogenetic relationship matrix that we fitted in the meta-analytic model did not explain a significant amount of variation. However, when splitting the data by classes of animals, we found not only that mammals and birds were most popular subjects in novel-object trials, but they also showed highest average repeatabilities. It seems plausible that highly visual organisms such as birds and many day-active mammals are particularly suitable for novel objects trials, (note, however, that the novel objects may also be detectable by odour). This finding is in agreement with the uneven representation of taxonomic classes observed by Rosenthal, Gertler, Hamilton, Prasad, and Andrade (2017). Our view on the consistency of responses to novel objects is thus strongly dominated by these two groups of vertebrates.

Overall, we found only minor publication bias in the published record. Furthermore, we found no difference in the magnitude of repeatability estimates between studies that focus on novel-object responses as the sole behaviour as compared to the large number of studies that combined multiple testing paradigms to evaluate personality dimensions. The robustly large amount of inter-individual variation in response to novel objects reliably produces significant repeatabilities, such that there is little scope for selective reporting and thus publication bias (Forstmeier, Wagenmakers, & Parker, 2017). Encouragingly, both the average sample size of repeatedly tested individuals and the time interval between the test repeats have increased over the years. If this trend continues, it will reveal more reliable estimates and also more data on long-term behavioural consistency. In recent years, a typical samples size was around 50-60 individuals retested after about 1-2 months.

### Terminology

Besides the question of how well novel-object trials allow a quantification of consistent individual differences, another important question is which animal behaviour or personality axis they are best ascribed to – a problem of labelling. Many publications in our survey dive straight into labelling. Many published abstracts use terms like “boldness” and “exploration” without stating how these were defined. However, boldness and exploration are particularly ambiguous labels, since they are also often used for startle response and open field tests, respectively. Neophilia, or even more precisely, object-neophilia is a less ambiguous term that is almost exclusively used for behavioural in novel object trails. In any case, we suggest that abstracts, and not only methods sections, should clearly state the testing paradigms that were used in the quantification of individual differences. Mentioning the label is usually not conclusive enough.

Neophobia/neophilia might be seen as a component of either boldness or exploration. Réale et al. (2007) suggest that neophobia/neophilia are components of exploration, thus do not correspond to a distinct behavioural phenotype, and boldness-shyness are described as reaction to risky but not new situations. However, neophobia might also be interpreted as a behavioural response to a risky situation. It is often unclear if an animal will perceive a novel object as risky or neutral. If this was clear, one could draw a fine line between neophobia as response to risky novelty (more in line with boldness) and neophilia as response to neutral novelty (more in line with exploration). In most cases, how animals perceive the situation will not be known and a differentiation is thus ambiguous.

The most frequent terms used in order to describe the animals’ reactions to a novel object were neophobia/neophilia (48 studies), boldness-shyness (31 studies) and exploration (22 studies), whereas few studies were using multiple labels. An important difference between the above terms is the testing context used for their assessment. The terms neophobia-neophilia were used mostly when the novel object was placed in or close to a food source (thus amplifying the risk aspect). This seems suitable if animals are motivated to approach a food source, but are prevented from approach by ‘fear of the new’. When the novel object was placed in a neutral position (e.g. in the middle of the testing cage), the use of all three terms was distributed more evenly, which reflects that novelty might than be seen as a thing to be discovered and explored (thus amplifying the exploratory aspect) or as a risky situation that induces neophobia and thus requires boldness to approach.

It would be worthwhile to study if novel-object responses in a neutral context are better correlated with exploration and novel object responses close to food better with startle responses. However, we are not aware of any systematic review. For the time being, it seems best to label responses to novel objects as object neophilia/neophobia and to clearly specify if objects were placed close to a resource. Object neophobia/neophilia can then be interpreted as a component of boldness or exploration.

### Conclusions

We here evaluate current practices of novel object trials and estimate average effects when novel-object trials are used to estimate the magnitude of temporally consistent individual differences. We find that most studies replicate novel objects trials, that sample sizes have increased slightly over time and that there are more long-term assessments of behavioural consistencies. This illustrates overall good and improving research practice. Average effects tended to be even slightly larger than average behavioural consistencies across different testing paradigms, illustrating that the novel object paradigm is suitable for capturing individual consistencies to reveal differences among individuals. Almost all short-term studies used different novel objects for the test repeats, which seems important, while longterm studies use either the same or different novel objects. Our results suggest that the latter decision does not impact the results. While there is some variation in how behavioural traits are labelled, the most specific description would be object neophobia/neophilia that can be interpreted as a component of boldness or exploration. Because of overlap of labels with other testing paradigms, we suggest that abstracts of published papers specify the testing setup rather than referring only to labels.

## Acknowledgements

This research was funded by the German Research Foundation (DFG) as part of the SFB TRR 212 (NC3; funding INST 215/543-1, 396782608).

