## Supplement for "Novelty at second glance: A critical appraisal of the novel-object paradigm based on meta-analysis"

Abstract word count: 303

Word count main text: 6285

Declarations of interest: none

Acknowledgements: This research was funded by the German Research Foundation (DFG) as part of the SFB TRR 212 (NC3; funding INST 215/543-1, 396782608).

**SUPPLEMENTARY MATERIAL**

**Search query**

Web of Science

TS = (novel object*) OR
        (neophob* AND explorat*) OR
        (neophi* AND explorat*) OR
        ((bold* OR shy*) AND explorat*) OR
        (neophob*) OR
        (neophi*) OR
        (bold* OR shy*)PY = 1990-2020

WC = “Behavioral Sciences”

Language: (English)

Document types: (Article)

**Table S1. Exclusion criteria of title/abstract and full-text screening**

| **Exclusion criterion** | **Justification** | **Code name** |
| --- | --- | --- |
| Not empirical data | Papers do not present original empirical data to be meta-analysed. | No empirical study |
| Studies on human subjects | We were specifically interested in animal studies where methodological approaches are very different from studies on humans. | No non-human animals |
| Studies with no novel stimuli | When stimuli are either familiar (already existing in the animal’s environment) or irrelevant to the novel object test (e.g. maze), they are not considered novel-object trials. | No novel object test |
| Novel object test was conducted in order to assess other attributes but not any behaviour or personality traits | Several papers assess cognition, but cognitive response might be very different from the behavioural response that we are interested here. | No behavioural study |
| Novel object trials used to test for differences between treatment groups or clonal lines | If only clonal lines are compared, then the correlation is more within lines than within individuals. | No individual-based study |
| Novel object test was conducted only once per individual | At least two trials of novel objects are needed to calculate the within-individual correlation. | No replication |
| Correlation and/or repeatability measures were not reported | The effect size of interest is needed to be meta-analysed. | No effect size |

**Table S2. Descriptive measures of the dataset 1990-2020**

|  |  | **Number of effect sizes**  **N = 254** | **Number of studies**  **N = 113** |
| --- | --- | --- | --- |
| **Use of different novel objects** | |  |  |
|  | Yes | 156 | 74 |
|  | No | 98 | 33 |
|  | Both | - | 6 |
| **Domestication level and testing context** | |  |  |
|  | Domestic animals tested in artificial environment | 29 | 11 |
|  | Lab-reared animals tested in artificial environment | 110 | 54 |
|  | Captive wild animals tested in artificial environment | 77 | 29 |
|  | Wild animals tested in natural environment | 38 | 19 |
| **Testing context** | |  |  |
|  | Novel object in neutral position | 185 | 75 |
|  | Novel object close/inside nest | 15 | 8 |
|  | Novel object close/inside feeder | 54 | 30 |

**Table S3. Input variables of the meta-analytic model**

| **Variable** | **Type** | **Code** | **Levels** | **Explanation** |
| --- | --- | --- | --- | --- |
| Study ID | Random | StudyID | Categorical: 109 studies | Unique identifier for each study |
| Effect Size ID | Random | EffectSizeID | Categorical: 201 effect sizes | Unique identifier for each effect size |
| Species | Random | Species | Categorical: 67 species | Unique species names |
| Sample Size | Fixed | Sample | Numerical | Number of individuals tested repeatedly |
| Time | Fixed | Time | Numerical | Time interval between two trials |
| Novelty | Fixed | Novelty | Binary: 0: No, 1: Yes | Same or different objects used in repeated trials |
| Domestication status | Fixed | Domestication | Categorical: 1: Domestic animals tested in lab, 2: Lab-reared animals tested in lab, 3: Wild-caught animals tested in lab, 4: Wild animals tested in field | Domestication of species might affect the explorative behaviour of individuals. The place of testing might also has an impact on behavioural responses |
| GLMM | Fixed | GLMM | Binary: 0: No non-Gaussian linear model, 1: non-Gaussian linear model | Non-Gaussian linear models (e.g. Poisson and Binomial models) often lead to lower repeatabilities |
| Response type | Fixed | Response | Categorical: Single (behaviour), Composite (of multiple behaviours), Average (across multiple trials) | Composite measures and in particular averaged behaviours are expected to yield higher repeatabilities, because measurement error is reduced |
| Multiple Assays | Fixed | MultiAssaysYN | Binary: 0: Only novel object, 1: Multiple behavioural tests | Whether the novel object test was the only behavioural test of the study might influence the bias towards reporting only statistically significant results |
| Context | Fixed | Context | Categorical: Food, Nest, Neutral | Whether the novel object was placed next to a food item, next to or inside a nest or in a neutral spot might affect the estimate of individual differences |

**Table S4. Number of studies corresponding to combinations of terminology and testing contexts.**

| **Term** | **Neutral** | **Food** | **Nest** | **Total**  **(unique)** |
| --- | --- | --- | --- | --- |
| **Neophobia/Neophilia** | **24** | **24** | **4** | **48** |
| Neophobia | 14 | 24 | 4 |  |
| Neophilia | 10 | 0 | 0 |  |
| **Boldness-Shyness** | **26** | **5** | **1** | **31** |
| Boldness | 25 | 5 | 1 |  |
| Shyness | 1 | 0 | 0 |  |
| **Exploration** | **18** | **2** | **2** | **22** |
| **Other terms** | **3** | **0** | **2** | **5** |
| Activity | 1 | 0 | 0 |  |
| Approach-avoidance | 2 | 0 | 2 |  |
| Risk-responsiveness | 2 | 0 | 0 |  |
| Fearfulness | 5 | 0 | 0 |  |
| **No specific terms** | **8** | **0** | **0** | **8** |
| **Total (unique studies)** | **80** | **30** | **8** |  |

**Table S5. Number of studies per terminology used and moderators tested.**

|  |  | **Neophobia/Neophilia** | **Boldness-Shyness** | **Exploration** |
| --- | --- | --- | --- | --- |
| **Clade** | |  |  |  |
|  | Birds | 32 | 7 | 9 |
|  | Mammals | 2 | 13 | 5 |
|  | Fish | 1 | 5 | 2 |
|  | Insects | 0 | 4 | 0 |
|  | Reptiles | 2 | 0 | 0 |
| **Context** | |  |  |  |
|  | Food | 19 | 4 | 1 |
|  | Nest | 3 | 1 | 1 |
|  | Neutral | 17 | 24 | 14 |
| **Novelty** | |  |  |  |
|  | Different objects | 32 | 21 | 10 |
|  | Same object | 6 | 9 | 7 |
| **Domestication status** | |  |  |  |
|  | Domesticated | 5 | 2 | 2 |
|  | Lab-reared | 10 | 20 | 5 |
|  | Wild-caught | 18 | 3 | 5 |
|  | Wild | 5 | 4 | 4 |

**Table S6. Publications included in the final dataset.**

| **Study ID** | **Authors (Year)** | **Title** | **Journal** | **Species** |
| --- | --- | --- | --- | --- |
| 1 | Amy, Ung, Beguin, and Leboucher (2017) | Personality traits and behavioural profiles in the domestic canary are affected by sex and photoperiod | Ethology | *Serinus canaria* |
| 2 | An, Kriengwatana, Newman, MacDougall-Shackleton, and MacDougall-Shackleton (2011) | Social rank, neophobia and observational learning in black-capped chickadees | Behaviour | *Poecile atricapillus* |
| 3 | Arnold et al. (2016) | Individual variation in corticosterone and personality traits in the blue tit *Cyanistes caeruleus* | Behaviour | *Cyanistes caeruleus* |
| 4 | Basic, Winberg, Schjolden, Krogdahl, and Hoglund (2012) | Context-dependent responses to novelty in Rainbow trout (*Oncorhynchus mykiss*) selected for high and low post-stress cortisol responsiveness | Physiology and Behaviour | *Oncorhynchus mykiss* |
| 5 | Baxter-Gilbert, Riley, and Whiting (2019) | Bold New World: urbanization promotes an innate behavioral trait in a lizard | Behavioural Ecology Sociobiology | *Intellagama lesueurii* |
| 6 | Bibi et al. (2019) | Personality is associated with dominance in a social feeding context in the great tit | Behaviour | *Parus major* |
| 7 | Boogert, Reader, and Laland (2006) | The relation between social rank, neophobia and individual learning in starlings | Animal Behaviour | *Sturnus vulgaris* |
| 8 | Brust and Guenther (2017) | Stability of the guinea pigs personality - cognition - linkage over time | Behavioural Processes | *Cavia aperea* |
| 9 | Burns (2008) | The Validity of Three Tests of Temperament in Guppies (*Poecilia reticulata*) | Journal of Comparative Psychology | *Poecilia reticulata* |
| 10 | Carere, Drent, Privitera, Koolhaas, and Groothuis (2005) | Personalities in great tits, *Parus major*: stability and consistency | Animal Behaviour | *Parus major* |
| 11 | Collins, Hatch, Elliott, and Jacobs (2019) | Boldness, mate choice and reproductive success in *Rissa tridactyla* | Animal Behaviour | *Rissa tridactyla* |
| 12 | Coutant, Bagur, and Gilbert (2018) | Development of an observational quantitative temperament test in three common parrot species | Applied Animal Behaviour Science | *Amazona aestiva* |
| 13 | Damas-Moreira, Riley, Harris, and Whiting (2019) | Can behaviour explain invasion success? A comparison between sympatric invasive and native lizards | Animal Behaviour | *Podarcis sicula* |
| 14 | Dammhahn and Almeling (2012) | Is risk taking during foraging a personality trait? A field test for cross-context consistency in boldness | Animal Behaviour | *Microcebus murinus* |
| 15 | Dardenne, Ducatez, Cote, Poncin, and Stevens (2013) | Neophobia and social tolerance are related to breeding group size in a semi-colonial bird | Behavioural Ecology Sociobiology | *Hirundo rustica* |
| 16 | David, Auclair, and Cezilly (2011) | Personality predicts social dominance in female zebra finches, *Taeniopygia guttata*, in a feeding context | Animal Behaviour | *Taeniopygia guttata* |
| 17 | DeRango et al. (2019) | Intrinsic and maternal traits influence personality during early life in Galapagos sea lion, *Zalophus wollebaeki*, pups | Animal Behaviour | *Zalophus wollebaeki* |
| 18 | Devost, Jones, Cauchoix, Montreuil-Spencer, and Morand-Ferron (2016) | Personality does not predict social dominance in wild groups of black-capped chickadees | Animal Behaviour | *Poecile atricapillus* |
| 19 | Edwards, Burke, and Dugdale (2017) | Repeatable and heritable behavioural variation in a wild cooperative breeder | Behavioural Ecology | *Acrocephalus sechellensis* |
| 20 | Edwards, Dugdale, Richardson, Komdeur, and Burke (2018) | Extra-pair parentage and personality in a cooperatively breeding bird | Behavioural Ecology Sociobiology | *Acrocephalus sechellensis* |
| 21 | Ensminger and Westneat (2012) | Individual and Sex Differences in Habituation and Neophobia in House Sparrows (*Passer domesticus*) | Ethology | *Passer domesticus* |
| 22 | Exnerova et al. (2015) | Different reactions to aposematic prey in 2 geographically distant populations of great tits | Behavioral Ecology | *Parus major* |
| 23 | Farrell, Weaver, An, and MacDougall-Shackleton (2012) | Song bout length is indicative of spatial learning in European starlings | Behavioral Ecology | *Sturnus vulgaris* |
| 24 | Finkemeier, Trillmich, and Guenther (2016) | Match-Mismatch experiments using photoperiod expose developmental plasticity of personality traits | Ethology | *Cavia aperea* |
| 25 | Fox and Millam (2010) | The Use of Ratings and Direct Behavioural Observation to Measure Temperament Traits in Cockatiels (*Nymphicus hollandicus*) | Ethology | *Nymphicus hollandicus* |
| 26 | Frost et al. (2013) | Environmental change alters personality in the rainbow trout, *Oncorhynchus mykiss* | Animal Behaviour | *Oncorhynchus mykiss* |
| 27 | Funghi, Leitão, Ferreira, Mota, and Cardoso (2015) | Social Dominance in a Gregarious Bird is Related to Body Size But not to Standard Personality Assays | Ethology | *Estrilda astrild* |
| 28 | Gabriel and Black (2010) | Behavioural syndromes in Steller’s jays: the role of time frames in the assessment of behavioural traits | Animal Behaviour | *Cyanocitta stelleri* |
| 29 | Garamszegi et al. (2015) | Among-year variation in the repeatability, within- and between-individual, and phenotypic correlations of behaviors in a natural population | Behavioral Ecology and Sociobiology | *Ficedula albicollis* |
| 30 | Garamszegi et al. (2012) | Corticosterone, Avoidance of Novelty, Risk-Taking and Aggression in a Wild Bird: No Evidence for Pleiotropic Effects | Ethology | *Ficedula albicollis* |
| 31 | Grace and Anderson (2014) | Personality correlates with contextual plasticity in a free-living, long-lived seabird | Behaviour | *Sula granti* |
| 32 | Greenberg and Holekamp (2017) | Human disturbance affects personality development in a wild carnivore | Animal Behaviour | *Crocuta crocuta* |
| 33 | Greggor, Jolles, Thornton, and Clayton (2016) | Seasonal changes in neophobia and its consistency in rooks: the effect of novelty type and dominance position | Animal Behaviour | *Corvus frugilegus* |
| 34 | Greggor, Masuda, Flanagan, and Swaisgood (2020) | Age-related patterns of neophobia in an endangered island crow: implications for conservation and natural history | Animal Behaviour | *Corvus hawaiiensis* |
| 35 | Grindstaff, Hunsaker, and Cox (2012) | Maternal and developmental immune challenges alter behavior and learning ability of offspring | Hormones and behavior | *Taeniopygia guttata* |
| 36 | Guenther and Brust (2017) | Individual consistency in multiple cognitive performance: behavioural versus cognitive syndromes | Animal Behaviour | *Cavia aperea* |
| 37 | Guenther and Trillmich (2013) | Photoperiod influences the behavioral and physiological phenotype during ontogeny | Behavioral Ecology | *Cavia aperea* |
| 38 | Guenther, Brust, Dersen, and Trillmich (2014) | Learning and personality types are related in Cavies (*Cavia aperea*) | Journal of Comparative Psychology | *Cavia aperea* |
| 39 | Guenther, Finkemeier, and Trillmich (2014) | The ontogeny of personality in the wild guinea pig | Animal Behaviour | *Cavia aperea* |
| 40 | Guenther, Groothuis, Krüger, and Goerlich-Jansson (2018) | Cortisol during adolescence organises personality traits and behavioural syndromes | Hormones and Behaviour | *Cavia aperea* |
| 41 | Guido, Biondi, Vasallo, and Muzio (2017) | Neophobia is negatively related to reversal learning ability in females of a generalist bird of prey, the Chimango Caracara, *Milvago chimango* | Animal Cognition | *Milvago chimango* |
| 42 | Gyuris, Feró, and Barta (2012) | Personality traits across ontogeny in firebugs, *Pyrrhocoris apterus* | Animal Behaviour | *Pyrrhocoris apterus* |
| 43 | Haage, Bergvall, Maran, Kiik, and Angerbjörn (2013) | Situation and context impacts the expression of personality: The influence of breeding season and test context | Behavioural Processes | *Mustela lutreola* |
| 44 | Hebert, Lavin, Marks, and Dzieweczynski (2014) | The effects of 17a-ethinyloestradiol on boldness and its relationship to decision making in male Siamese fighting fish | Animal Behaviour | *Betta splendens* |
| 45 | Herborn et al. (2010) | Personality in captivity reflects personality in the wild | Animal Behaviour | *Cyanistes caeruleus* |
| 46 | Hirata and Arimoto (2018) | Novel object response in beef cattle grazing a pasture as a group | Behavioural Processes | *Bos taurus* |
| 47 | Hopkins and Bennett (1994) | Handedness and Approach-Avoidance behavior in Chimpanzees (*Pan*) | Journal of Experimental Psychology | *Pan troglodytes* |
| 48 | Jäger, Schradin, Pillay, and Rimbach (2017) | Active and explorative individuals are often restless and excluded from studies measuring resting metabolic rate: Do alternative metabolic rate measures offer a solution? | Physiology & Behaviour | *Rhabdomys pumilio* |
| 49 | Janczak, Pedersen, and Bakken (2003) | Aggression, fearfulness and coping styles in female pigs | Applied Animal Behaviour Science | *Sus domesticus* |
| 50 | Johnson et al. (2015) | Genetic Influences on Response to Novel Objects and Dimensions of Personality in *Papio* Baboons | Behavior Genetics | *Papio* |
| 51 | Jolles, Ostojic, and Clayton (2013) | Dominance, pair bonds and boldness determine social-foraging tactics in rooks, *Corvus frugilegus* | Animal Behaviour | *Corvus frugilegus* |
| 52 | Jolly, Webb, Gillespie, Hughes, and Phillips (2019) | Bias averted: personality may not influence trappability | Behavioural Ecology Sociobiology | *Melomys burtoni* |
| 53 | Kerman, Miller, and Sewall (2018) | The effect of social context on measures of boldness: Zebra finches (*Taeniopygia guttata*) are bolder when housed individually | Behavioural Processes | *Taeniopygia guttata* |
| 54 | Krams et al. (2014) | Sex-Specific associations between nest defence, exploration and breathing rate in breeding pied flycatchers | Ethology | *Ficedula hypoleuca* |
| 55 | Krause, Krüger, and Schielzeth (2017) | Long-term effects of early nutrition and environmental matching on developmental and personality traits in zebra finches | Animal Behaviour | *Taeniopygia guttata* |
| 56 | Krebs, Linnenbrink, and Guenther (2019) | Validating standardised personality tests under semi-natural conditions in wild house mice (*Mus musculus domesticus*) | Ethology | *Mus domesticus* |
| 57 | Kurvers et al. (2009) | Personality differences explain leadership in barnacle geese | Animal Behaviour | *Branta leucopsis* |
| 58 | Kurvers, de Hoog, van Wieren, Ydenberg, and Prins (2012) | No evidence for negative frequency-dependent feeding performance in relation to personality | Behavioural Ecology | *Branta leucopsis* |
| 59 | Le Vin, Mable, Taborsky, Heg, and Arnold (2011) | Individual variation in helping in a cooperative breeder: relatedness versus behavioural type | Animal Behaviour | *Neolamprologus pulcher* |
| 60 | Lermite, Peneaux, and Griffin (2017) | Personality and problem-solving in common mynas (*Acridotheres tristis*) | Behavioural Processes | *Acridotheres tristis* |
| 61 | Malmkvist and Hansen (2002) | Generalization of fear in farm mink, *Mustela vison*, genetically selected for behaviour towards humans | Animal Behaviour | *Neovison vison* |
| 62 | Martin-Wintle et al. (2017) | Do opposites attract? Effects of personality matching in breeding pairs of captive giant pandas on reproductive success | Biological Conservation | *Ailuropoda melanoleuca* |
| 63 | Mazza et al. (2019) | Coping with style: individual differences in responses to environmental variation | Behavioural Ecology Sociobiology | *Myodes glareolus* |
| 64 | Mazza, Eccard, Zaccaroni, Jacob, and Dammhahn (2018) | The fast and the flexible: cognitive style drives individual variation in cognition in a small mammal | Animal Behaviour | *Myodes glareolus* |
| 65 | McCune, Jablonski, Lee, and Ha (2018) | Evidence for personality conformity, not social niche specialization in social jays | Behavioral Ecology | *Aphelocoma wollweberi* |
| 66 | Meagher, von Keyserlingk, Atkinson, and Weary (2016) | Inconsistency in dairy calves’ responses to tests of fearfulness | Applied Animal Behaviour Science | *Bos taurus* |
| 67 | Medina-Garcia, Jawor, and Wright (2017) | Cognition, personality, and stress in budgerigars, *Melopsittacus undulatus* | Behavioral Ecology | *Melopsittacus undulatus* |
| 68 | Meehan and Mench (2002) | Environmental enrichment affects the fear and exploratory responses to novelty of young Amazon parrots. | Applied Animal Behaviour Science | *Amazona amazonica* |
| 69 | Mettke-Hofmann (2012) | Head Colour and Age Relate to Personality Traits in Gouldian Finches | Ethology | *Erythrura gouldiae* |
| 70 | Mettke-Hofmann, Ebert, Schmidt, Steiger, and Stieb (2005) | Personality traits in resident and migratory warbler species | Behaviour | *Sylvia melanocephala* |
| 71 | Michelena, Sibbald, Erhard, and McLeod (2009) | Effects of group size and personality on social foraging: the distribution of sheep across patches | Behavioral Ecology | *Ovis aries* |
| 72 | Miller, Garner, and Mench (2005) | The test-retest reliability of four behavioural tests of fearfulness for quail: a critical evaluation | Applied Animal Behaviour Science | *Coturnix coturnix japonica* |
| 73 | Miller, Garner, and Mench (2006) | Is fearfulness a trait that can be measured with behavioural tests? A validation of four fear tests for Japanese quail | Animal Behaviour | *Coturnix coturnix japonica* |
| 74 | Moldoff and Westneat (2017) | Foraging sparrows exhibit individual differences but not a syndrome when responding to multiple kinds of novelty | Behavioral Ecology | *Passer domesticus* |
| 75 | Monestier et al. (2017) | Neophobia is linked to behavioural and haematological indicators of stress in captive roe deer | Animal Behaviour | *Capreolus capreolus* |
| 76 | Morinay, Daniel, Gustafsson, and Doligez (2019) | No evidence for behavioural syndrome and genetic basis for three personality traits in a wild bird population | Animal Behaviour | *Ficedula albicollis* |
| 77 | Noer, Needham, Wiese, Balsby, and Dabelsteen (2016) | Personality matters: Consistency of inter-individual variation in shyness-boldness across non-breeding and pre-breeding season despite a fall in general shyness levels in farmed American mink (*Neovison vison*) | Applied Animal Behaviour Science | *Neovison vison* |
| 78 | Overington, Cauchard, Cote, and Lefebvre (2011) | Innovative foraging behaviour in birds: what characterizes an innovator? | Behavioural Processes | *Quiscalus lugubris* |
| 79 | Pedersen (1994) | Long-term effects of different handling procedures on behavioural, physiological and production-related parameters in silver foxes | Applied Animal Behaviour Science | *Vulpes vulpes* |
| 80 | Perals, Griffin, Bartomeus, and Sol (2017) | Revisiting the open-field test: what does it really tell us about animal personality? | Animal Behaviour | *Acridotheres tristis* |
| 81 | Pogány et al. (2018) | Personality assortative female mating preferences in a songbird | Behaviour | *Taeniopygia guttata* |
| 82 | Rangassamy et al. (2016) | Personality modulates proportions of CD4+ regulatory and effector T cells in response to socially induced stress in a rodent of wild origin. | Physiology and Behaviour | *Mus spicilegus* |
| 83 | Rockwell, Gabriel, and Black (2012) | Bolder, older, and selective: factors of individual-specific foraging behaviors in Steller’s jays | Behavioural Ecology | *Cyanocitta stelleri* |
| 84 | Rohrer and Ferkin (2020) | Long-term repeatability and stability of three personality traits in meadow voles | Ethology | *Microtus pennsylvanicus* |
| 85 | Ruuskanen and Laaksonen (2010) | Yolk hormones have sex-specific long-term effects on behavior in the pied flycatcher (*Ficedula hypoleuca*). | Hormones and Behavior | *Ficedula hypoleuca* |
| 86 | Schielzeth, Bolund, Kempenaers, and Forstmeier (2010) | Quantitative genetics and fitness consequences of neophilia in zebra finches | Behavioral Ecology | *Taeniopygia guttata* |
| 87 | Schürch and Heg (2010) | Life history and behavioral type in the highly social cichlid, *Neolamprologus pulcher* | Behavioural Ecology | *Neolamprologus pulcher* |
| 88 | Siviter et al. (2017) | The impact of egg incubation temperature on the personality of oviparous reptiles | Animal Cognition | *Pogona vitticeps* |
| 89 | Smith and Blumstein (2012) | Structural consistency of behavioural syndromes: does predator training lead to multi-contextual behavioural change? | Behaviour | *Poecilia reticulata* |
| 90 | Soha, Peters, Anderson, Searcy, and Nowicki (2019) | Performance on tests of cognitive ability is not repeatable across years in a songbird | Animal Behaviour | *Melospiza melodia* |
| 91 | Sol, Griffin, and Bartomeus (2012) | Consumer and motor innovation in the common myna: the role of motivation and emotional responses | Animal Behaviour | *Acridotheres tristis* |
| 92 | Spake, Gray, and Cassady (2012) | Relationship between back test and coping styles in pigs | Applied Animal Behaviour Science | *Sus domesticus* |
| 93 | Stöwe, Bugnyar, Heinrich, and Kotrschal (2006) | Effects of group size on approach to novel objects in ravens (*Corvus corax*) | Ethology | *Corvus corax* |
| 94 | Stöwe, Bugnyar, Loretto, et al. (2006) | Novel object exploration in ravens (*Corvus corax*): Effects of social relationships | Behavioural Processes | *Corvus corax* |
| 95 | Stuber et al. (2013) | Slow explorers take less risk: a problem of sampling bias in ecological studies | Behavioral Ecology | *Parus major* |
| 96 | Tan and Tan (2019) | Individual- and population-level personalities in a floriphilic katydid | Ethology | *Phaneroptera brevis* |
| 97 | Tobler and Sandell (2007) | Yolk testosterone modulates persistence of neophobic responses in adult zebra finches, *Taeniopygia guttata* | Hormones and Behavior | *Taeniopygia guttata* |
| 98 | Tremmel and Müller (2013) | Insect personality depends on environmental conditions | Behavioral Ecology | *Phaedon cochleariae* |
| 99 | Tremmel and Müller (2014) | Diet dependent experience and physiological state shape the behavior of a generalist herbivore | Physiology & Behavior | *Galeruca tanaceti* |
| 100 | Trompf and Brown (2014) | Personality affects learning and trade-offs between private and social information in guppies, *Poecilia reticulata* | Animal Behaviour | *Poecilia reticulata* |
| 101 | Valros, Pedersen, Poytakangas, and Jensen (2017) | Evaluating measures of exploratory behaviour in sows around farrowing and during lactation-A pilot study | Applied Animal Behaviour Science | *Sus scrofa domesticus* |
| 102 | Verbeek, Drent, and Wiepkema (1994) | Consistent individual differences in early exploratory behaviour of male great tits | Animal Behaviour | *Parus major* |
| 103 | Vernouillet and Kelly (2020) | Individual exploratory responses are not repeatable across time or context for four species of food storing corvid | Nature Scientific Reports | *Gymnorhinus cyanocephalus* |
| 104 | Vetter et al. (2016) | Shy is sometimes better: personality and juvenile body mass affect adult reproductive success in wild boards, *Sus scrofa* | Animal Behaviour | *Sus scrofa* |
| 105 | Vrublevska et al. (2015) | Personality and density affect nest defence and nest survival in the great tit | Acta Ethologica | *Parus major* |
| 106 | Williams, King, and Mettke-Hofmann (2012) | Colourful characters: head colour reflects personality in a social bird, the Gouldian finch, *Erythrura gouldiae* | Animal Behaviour | *Erythrura gouldiae* |
| 107 | Wilson and Stevens (2005) | Consistency in context-specific measures of shyness and boldness in rainbow trout, *Oncorhyncus mykiss* | Ethology | *Oncorhynchus mykiss* |
| 108 | Winter, Martins, Trovo, and Chapman (2016) | Different behaviour-body length correlations in two populations of juvenile three-spined stickleback (*Gasterosteus aculeatus*) | Behavioural Processes | *Gasterosteus aculeatus* |
| 109 | Yuen, Pillay, Heinrichs, Schoepf, and Schradin (2015) | Personality does not constrain social and behavioural flexibility in African striped mice | Behavioural Ecology Sociobiology | *Rhabdomys pumilio* |
| 110 | Yuen, Pillay, Heinrichs, Schoepf, and Schradin (2016) | Personality traits are consistent when measured in the field and in the laboratory in African striped mice (*Rhabdomys pumilio*) | Behavioural Ecology Sociobiology | *Rhabdomys pumilio* |
| 111 | Zidar et al. (2017) | A comparison of animal personality and coping styles in the red junglefowl | Animal Behaviour | *Gallus gallus* |
| 112 | Zidar et al. (2018) | The relationship between learning speed and personality is age- and task-dependent in red junglefowl | Behavioural Ecology Sociobiology | *Gallus gallus* |

**Fig. S1 Phylogenetic tree of species included in the meta-analysis**

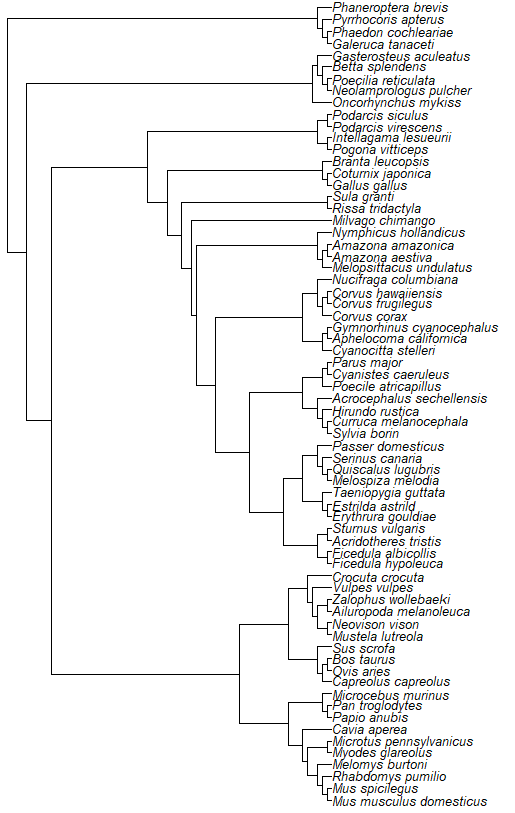

**Fig. S2. Influence diagnostics of effect sizes. Red points indicate outliers. X axes shows effect size ID.**

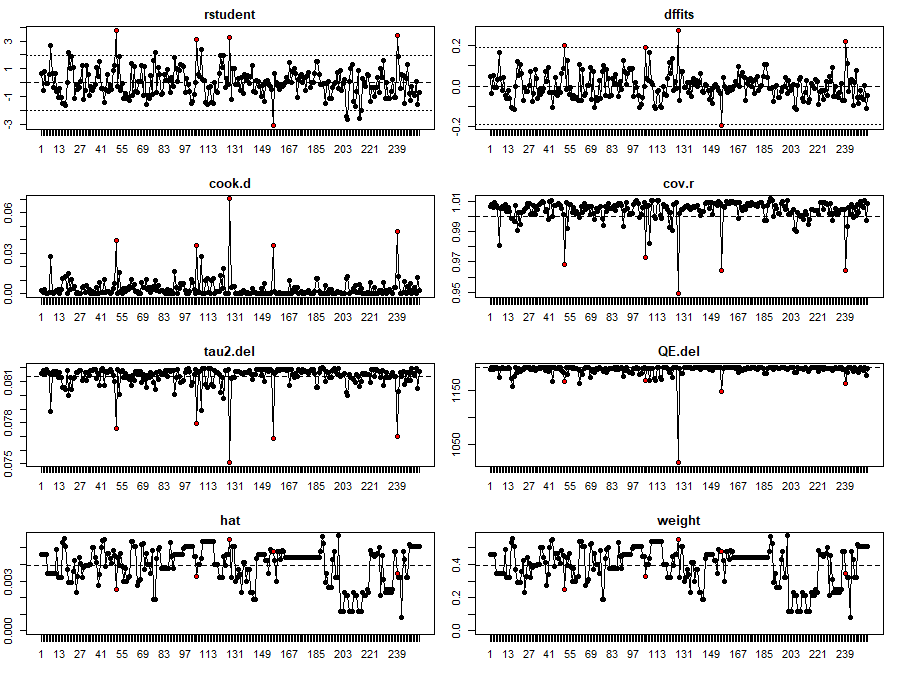

**Fig. S3. Heatmap showing the number of studies by response behaviour and label for the behavioural phenotypes.**

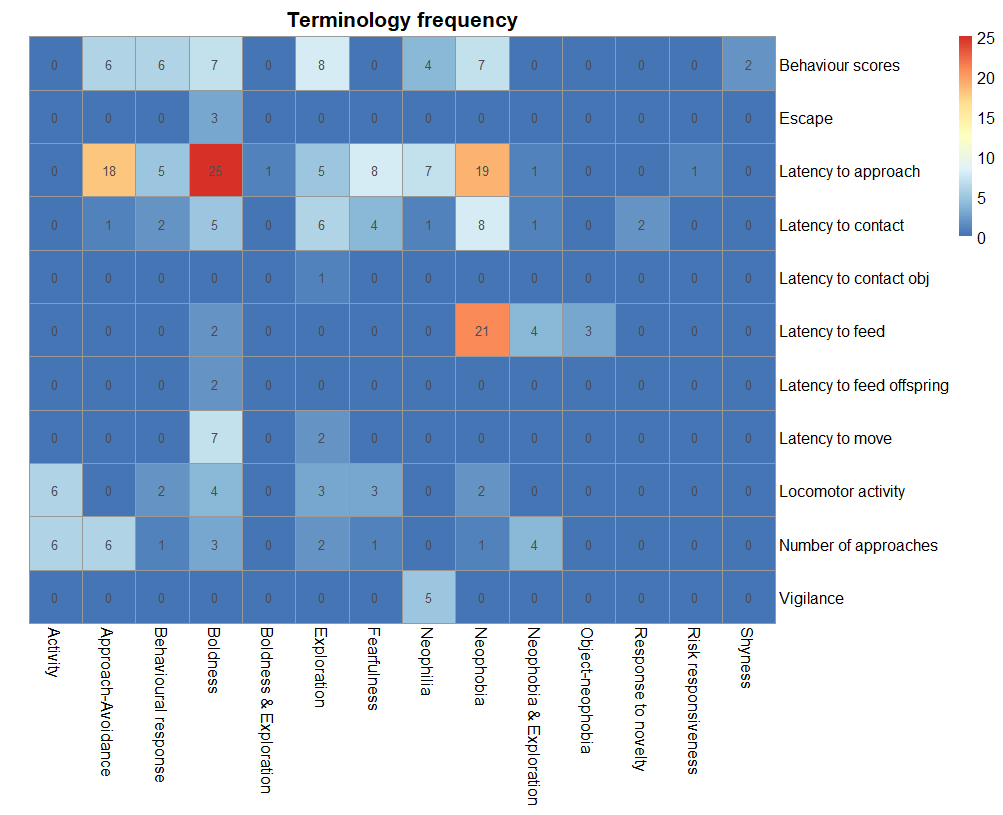

**Fig. S4. Number of studies per year. Different colours represent the use of same or different objects during repeated trials of novel object test.**

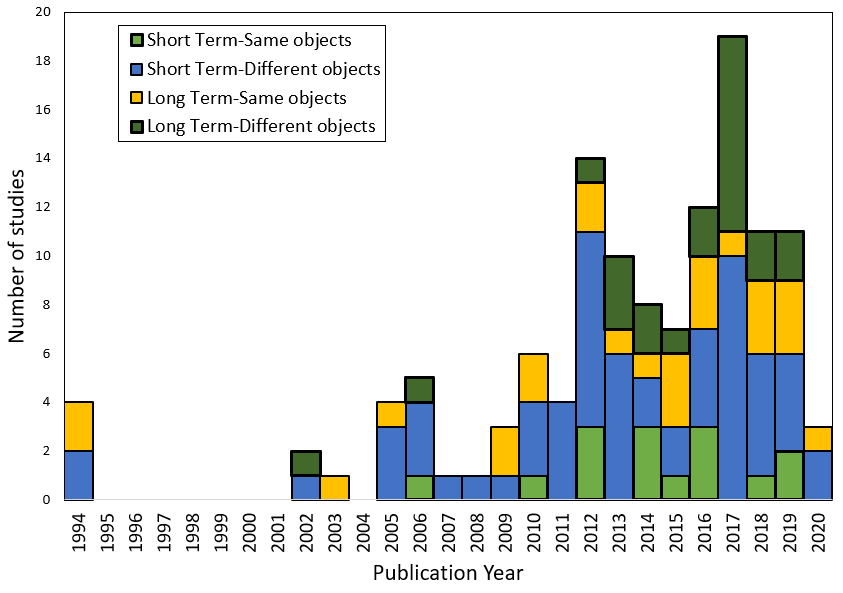
